# Molecular Dynamic Investigation of H5N1 Influenza Virus Dual H274Y-I222K Mutation Resistance to Peramivir

**DOI:** 10.1101/2022.03.08.483396

**Authors:** Ndumiso M. Buthelezi, Sphamandla E. Mtambo, Daniel G. Amoako, Anou M. Somboro, Elliasu Yakubu, Rene B. Khan, Ndumiso N. Mhlongo, Hezekiel M. Kumalo

**Author notes:** Correspondence (H.M.K), (D.G.A.).

## Abstract

In light of the rapid rise of an influenza pandemic, the constant genetic mutations of H5N1 influenza viruses pose a threat. Mutations at the sialic site are often responsible for multiple drug resistance. To design effective new inhibitors, it is necessary to undertake research into the mechanism of resistance of influenza viruses and their rapid mutations. The molecular dynamic simulation technique has been an instrumental tool in understanding how proteins function from an atomic perspective. A thorough investigation has not been conducted using molecular dynamics to examine the impact of these mutations (I222K, H274Y, and H274Y-I222K) on Peramivir. This study investigates the effects of I222K, H274Y, H274Y-I222K substitution on the neuraminidase–Peramivir complex and identifies responsible residues for complex conformations. The mutations caused distorted Peramivir orientation in the enzyme active site, which affected the inhibitor’s binding. In the presence of various mutations, interaction between protein and ligand became less thermodynamically favorable. We observed the following trend in binding free energy difference: WT<I222<H274Y<H274Y-I222K. As a result of the thermodynamic instability of the mutant complexes, Peramivir’s potency is reduced due to impaired binding interactions. Wild type complex displays thermodynamic stability and strong protein-ligand interactions due to their high total energy contributions and low residue flexibility. Based on the energy decomposition analysis, Arg117, Arg224, and Arg292 contributed the largest residual energy for the binding of Peramivir to wild type and mutants. These residues are thought to play a key role in the formation of the binding pocket between Peramivir and neuraminidase. This study provides a basis for investigating the effects of other mutations on Peramivir’s efficacy against the H5N1 virus.

## 1. Introduction

Influenza virus (H5N1) virus is a highly pathogenic avian influenza that has dramatically evolved since its discovery in Hong Kong, China, in May 1997 [1]. For the past 20 years, researchers have been puzzled and challenged by a virus that can poorly jump the species barrier into humans. Still, when it does, the outcome is catastrophically fatal. This has drawn awareness of efforts to develop plans for its control. Influenza viruses belong to the Orthomyxoviridae family with two types, A and B, which cause significant outbreaks in humans [2]. Influenza viruses have negative-sense, single-stranded, segmented RNA genomes and possess the membrane proteins, the hemagglutinin (HA) and the neuraminidase (NA). The HA protein mediates sialic acid-conjugated cell receptor recognition, receptor binding, and fusion-mediated entry into cells [3,4]. The NA protein facilitates the release of newly assembled virus particles by cleavage of sialic acid residues. Mutations commonly occur in the active site or proximity to the active region as shown in Figure 1 [5].

**Figure 1.**
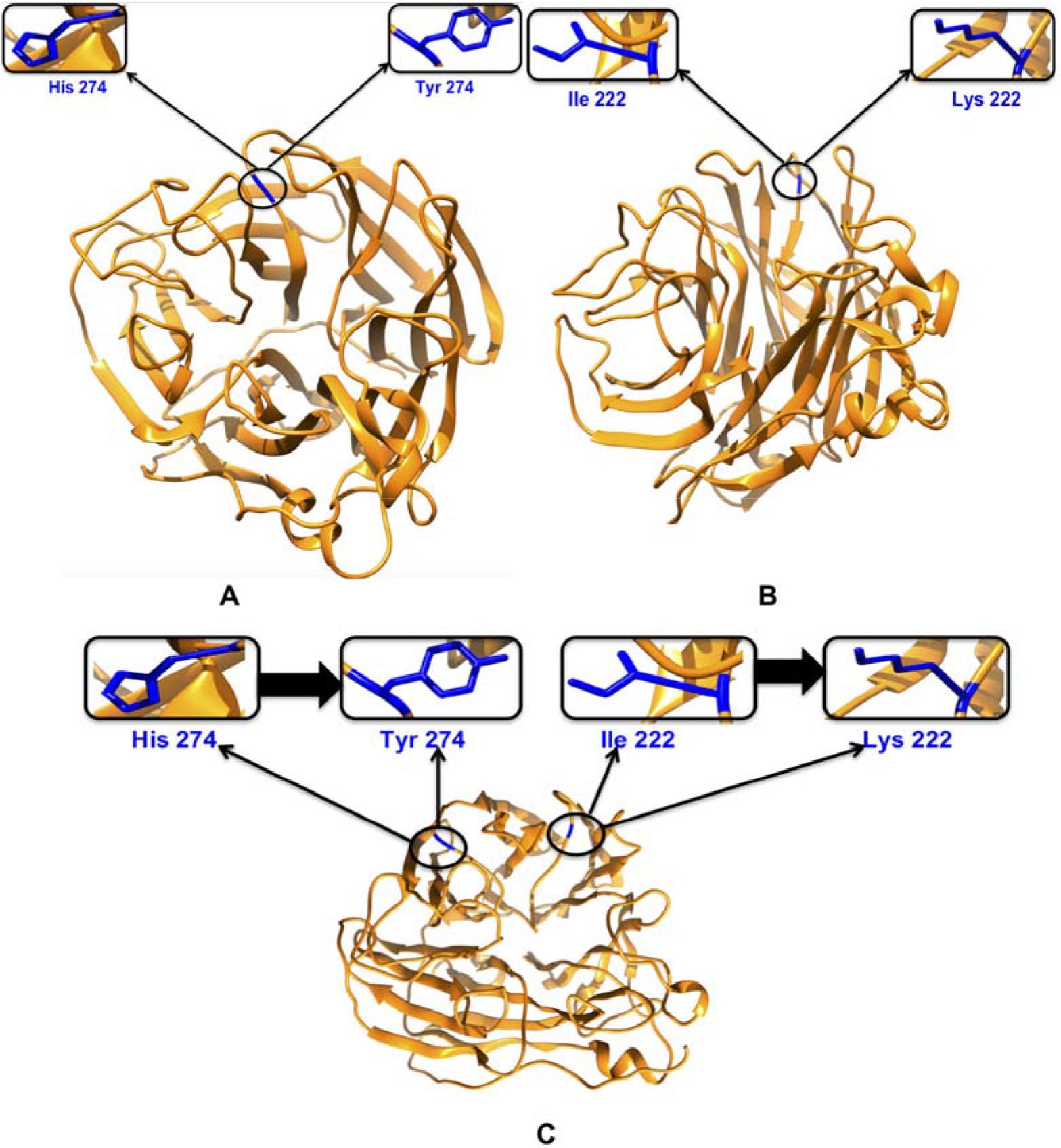
Depictions of the mutations carried out, H274Y (A), I222K (B), and H274Y-I222K (C).

**Figure 2.**
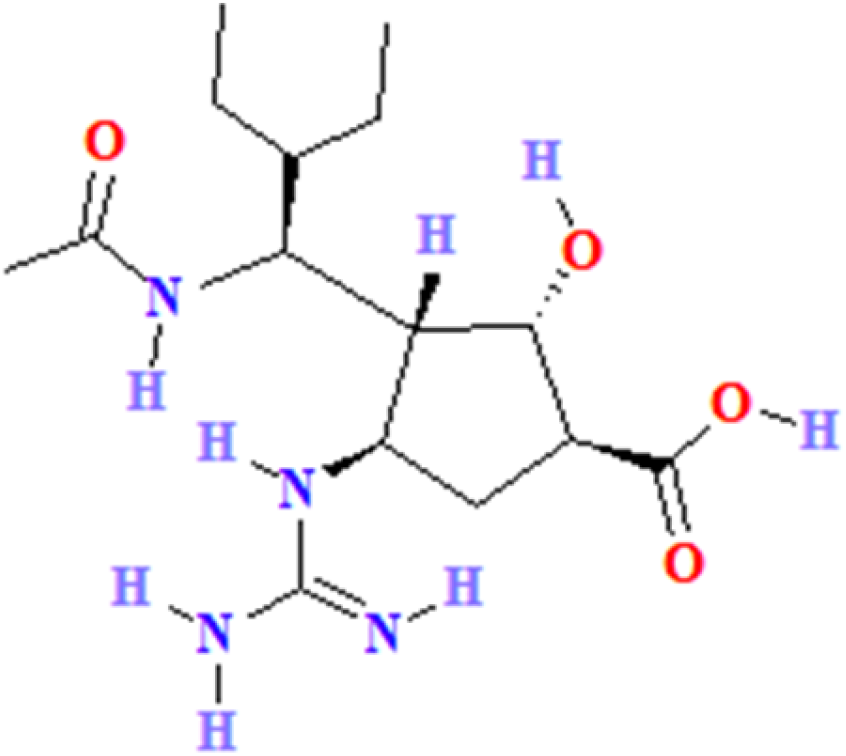
The chemical structure of Peramivir.

The constant genetic mutations of H5N1 influenza viruses pose a threat due to a rapid rise of influenza pandemic [1,6,7]. Mutations found in the sialic site are known culprits for multiple drug-resistant (MD-R) strain [8]. This has had an adverse effect on neuraminidase inhibitors potency. Neuraminidase inhibitors (NAIs) have helped fight the war against pandemic influenza worldwide. There are four neuraminidase inhibitors that are currently approved in various countries for the treatment of influenza, namely, Zanamivir, Oseltamivir, Laninamivir, and Peramivir are used to treat and prevent uncomplicated influenza virus infections [9].

The research into the deeper understanding of the mechanism of resistance of the influenza virus and its rapid mutations must be fast-tracked in order to design effective novel inhibitors. Molecular dynamic (MD) simulation has been an instrumental tool in gaining insight into atomic perspective of the complex nature of proteins as a function of time. Sequence identity between the H5N1, H1N1 (2009) and the conserved active site is 91.47%; therefore, the results from molecular dynamics simulation are applicable to both viruses [10].

Various methodologies, such as *in vitro* studies have been done to understand the impact of the mutations on Peramivir resistances to neuraminidase. The H274Y conversion increased the resistance of Peramivir to neuraminidase. In 2007, during the Euroasia influenza season, Karthick et al., pursued the first MD study on the natural mutation H274Y of H1N1 [11]. Similar studies based on the N294S and E119G by Wang and Zheng revealed that residue 274 is an active site residue that facilitates the binding of NA inhibitors [12]. The mutation of Histidine (His) to Tyrosine (Tyr) negatively impacted the binding pocket thus the pocket could not accommodate the steric bulkiness of the inhibitor. The dual mutation H274Y-I222K further increased resistance by 10- to 90-fold [13]. The mutation at position 222 from Isoleucine (Iso) to Lysine (Lys) was also found to disturb the framework stability.

The impact of these mutation I222K, H274Y and H274Y-I222K on Peramivir has not been thoroughly using molecular dynamics. Reports of the emergence of drug resistance make the development of new anti-influenza molecules a priority. Therefore, mapping the drug interactions of Peramivir (Figure 1) against the different mutations will help with the design of novel inhibitors. The insufficient computational molecular dynamics of the resistances of Peramivir against neuraminidase H5N1 mutations, I222K and H274Y prompted us perform a comprehensive analysis.

## 2. Results and Discussion

### 2.1. Root-Mean-Square Deviations (RMSD)

A graphical monitor of the convergence of the systems was monitored using the RMSD and potential energy plots prior to MD analysis [14]. All the systems converge at approximately 5, 000 ps for both the RMSD and possible plot. The RMSD is a similarity measure applied to estimate the deviation in the degree of protein stability. A high RMSD implies increased mobility of backbone atoms indicative of structural instability, while a low RMSD value signifies a decreased mobility of the backbone C-α atoms and protein stability. Thus, a high RMSD signifies a disruption in enzyme structural framework. In Figure 3, the H274Y mutant showed a higher RMSD when compared to the wild type. H274Y mutant is a distortion of the structural framework hence the decrease in protein stability.

**Figure 3.**
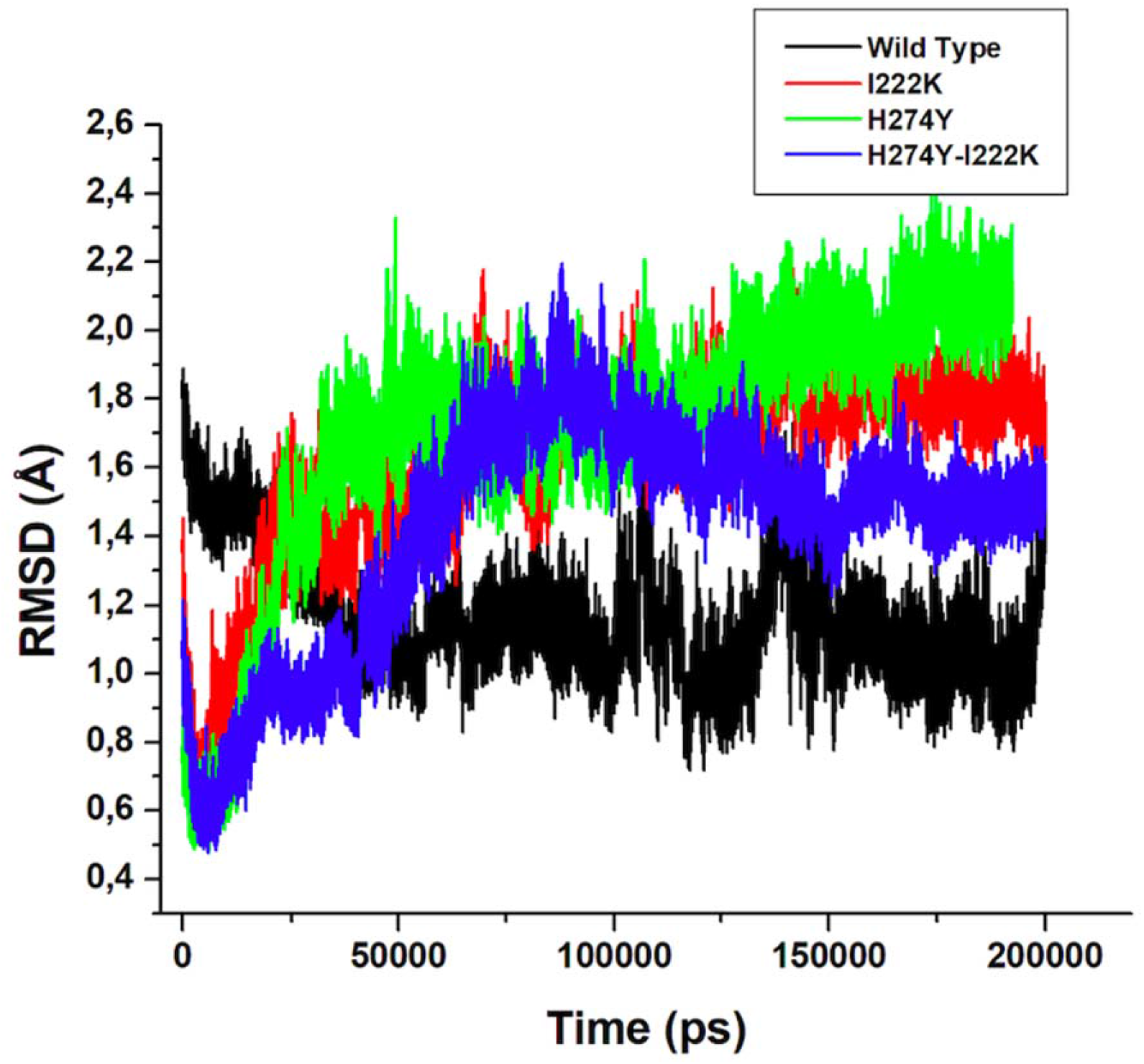
RMSD plot of C-α atoms of the wild type, H274Y, I222K and H274Y-I222K mutations.

### 2.2. Root-Mean-Square Fluctuation (RMSF)

In order to gain insight into the atomic fluctuations an observation between the residues or atoms present in a molecule therefore applied to give insight into the flexible regions of the protein during the simulation [15]. A distinction is made between high and low fluctuations where higher fluctuations indicate increased flexibility and or vice versa. In Figure 4, an illustration of the amino acid flexibility for all wild type, I222K, H274Y and H274Y-I222K systems is shown. An observation made was that the systems shared a similar trend in residual fluctuations. However, the H274Y-I222K, H274Y and I222K mutants shows a higher degree of flexibility in the region 59–73, 150–155, 163–169, 187–194, 257–263, 270–280, 302–307, 346–356 and 368–384 when compared to wild type.

**Figure 4.**
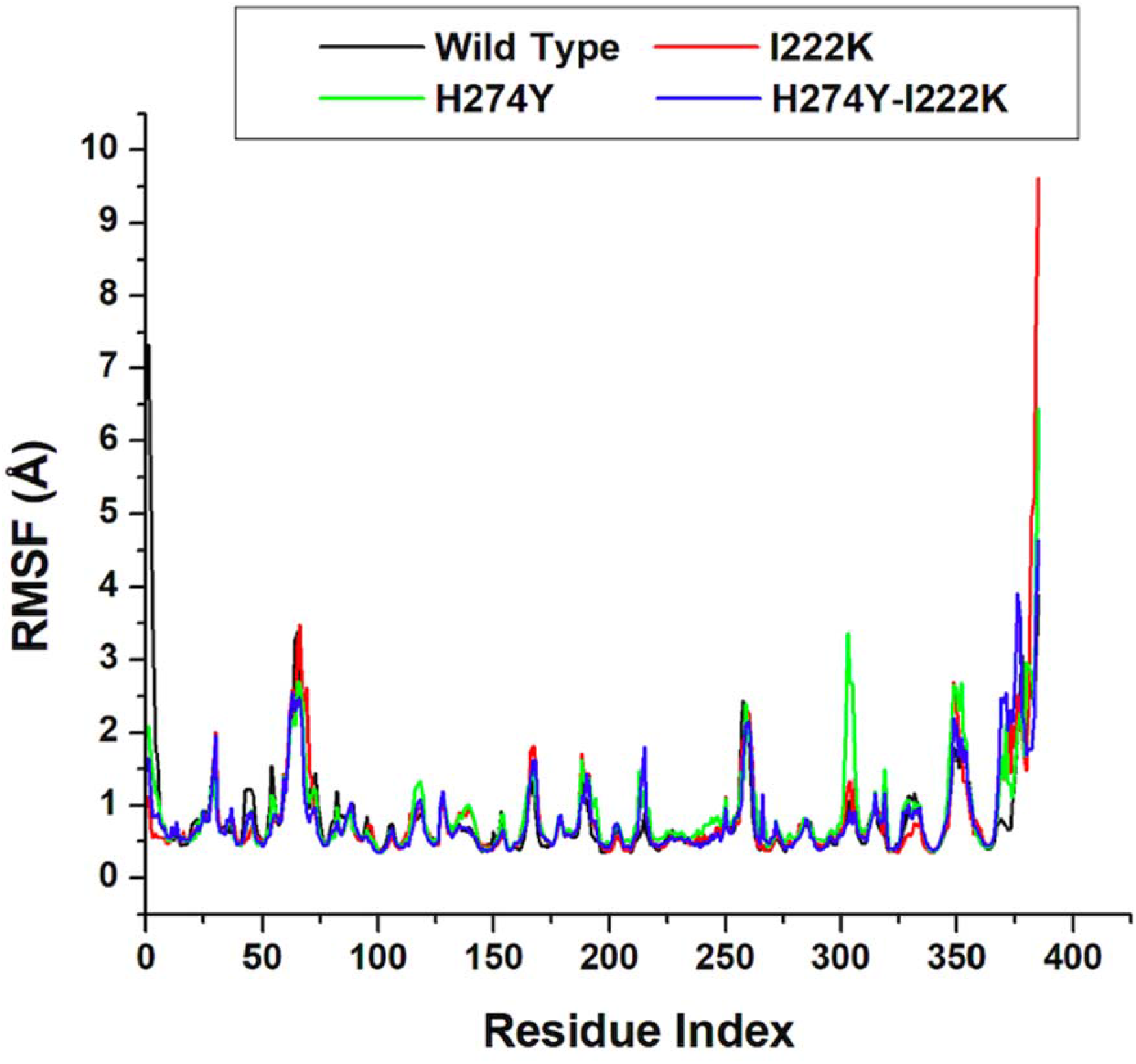
Residue-based average C-α fluctuations of the wild type, I222K, H274Yand H274Y-I222K mutations.

The other significant note is that the mutants showed a higher fluctuation degree in the region 150–155 where the ASP151 is located. The Asp151 loop plays a significant role in substrate binding due to its open and closed cavity of the binding site. Whilst the wild type displayed lower fluctuations for similar amino acid residues. The residue region 260–320 containing the compensatory mutation did not demonstrate significant variations in fluctuations. The role of residue 222 appeared to shift from a structural one to a functional one. The Lys222 mutation contains a second amino group that acts as an electrophile at physiological pH; this prompts interaction with the active site residues.

The change from a hydrophobic residue, Histidine, to the aromatic Tyrosine at position 274, which contains an ionizable phenolic group destabilized, introduced instability in a previously predominant hydrophobic region. Therefore, it is plausible to assume that the presence of mutation I222K in system H274Y-I222K restricts interaction with the external solvent, thus minimizing flexibility throughout the protein. Comparing the single and double mutations, the RMSF shows that both systems indicate no difference between the two. Therefore, this means that the conformation of the protein is not disturbed.

### 2.3. Radius of Gyration (RoG)

System compactness along the simulation was computed to gain insight into the tertiary protein structure and provide insight into complex stability and changes of the systems along the MD simulation [16,17].

To measure the overall protein dimensions of H5N1 systems, RoG was plotted and analyzed in Figure 5. From Figure 5, it is evident that the systems share close resemblance, which is explained by the similar arrangement of the amino acids in the secondary and tertiary structures. H274Y system showed a gradual increment. Comparing wild type and I222K indicates that the mutation is a more compact system than the wild. The double mutation, H274Y-I222K, showed more flexibility when compared to the single mutation H274Y. An assumption is that the double mutation attempts to preserve the active site by interfering with the Peramivir binding. Upon closer analysis of similar complex compactness is observed between the H274Y and H274Y-I222K.

**Figure 5.**
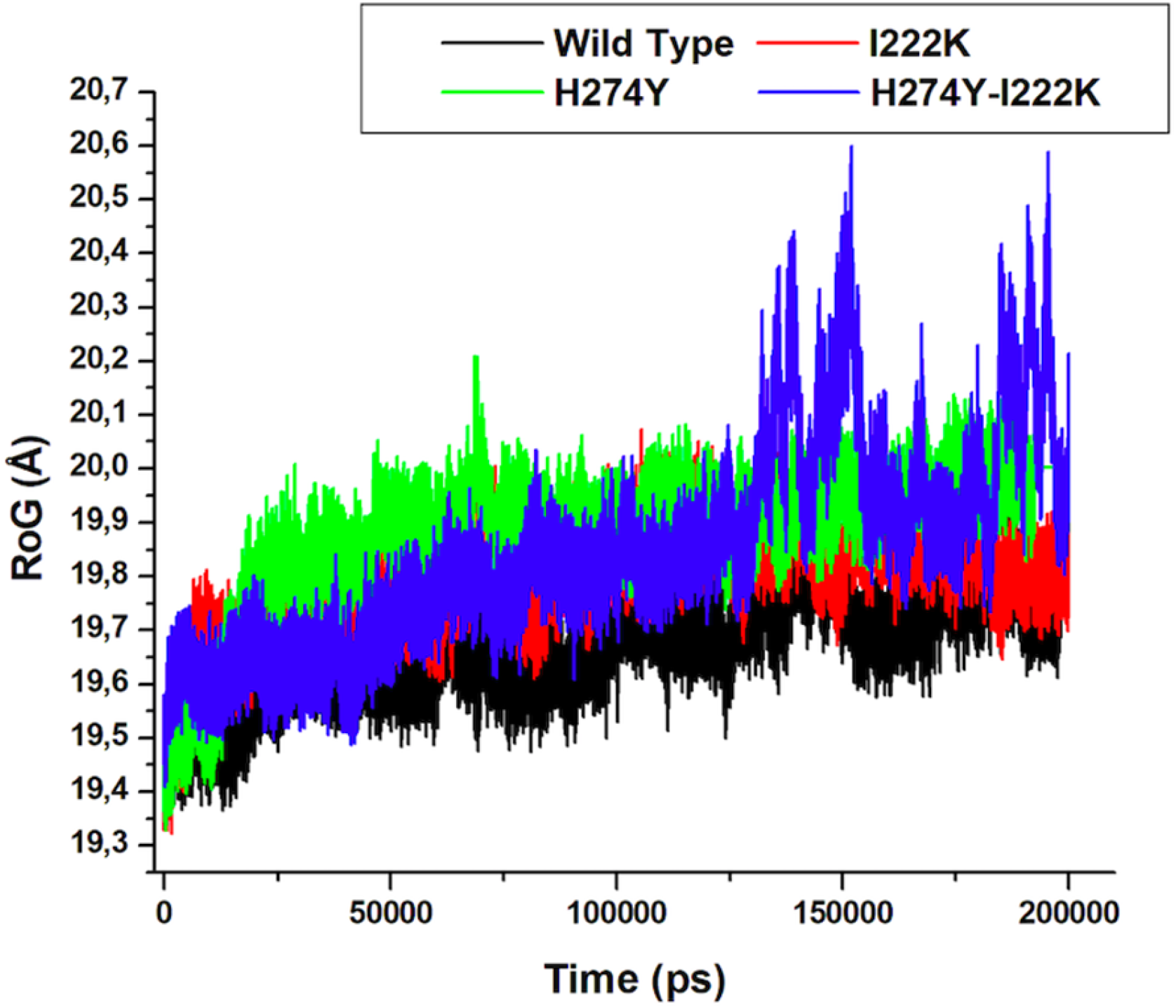
Radius of gyration of the wild type, I222K, H274Y and H274Y-I222K mutant systems measured over 200,000 ps MD simulation.

### 2.4. MM/GBSA Binding Free Energy Calculation

Thermodynamic calculations were carried out for the wild type, H274Y, I222K and H274Y-I222K mutations. Thermodynamic calculations were used as a tool that provides a thorough insight into the impact(s) of the mutation(s) on the Peramivir binding affinity in the H5N1 enzyme cavity. The relative binding free energy of the protein-ligand complexes was averaged over 1000 snapshots extracted from the 100 000 ps MD trajectories using the MM/GBSA technique. Table 1 illustrates the binding affinity profiles of Peramivir bound to the wild type, H274Y, I222K and H274Y-I222K mutation(s). All molecular mechanics and solvation energy components were calculated using the MM/GBSA approach over a 200 ns MD trajectory, as listed in Table 1.

**Table 1.** MM/GBSA based binding free energy profile of Peramivir bound to wild type, H274Y, I222K and H274Y-I222K.

| Complexes | $\Delta G_{\text{bind}}$ | $\Delta E_{\text{ele}}$ | $\Delta E_{\text{vdw}}$ | $\Delta E_{\text{gas}}$ | $\Delta G_{\text{sol}}$ |
| --- | --- | --- | --- | --- | --- |
| Wild type | $-29.96 \pm 4.88$ | $-101.50 \pm 15.85$ | $-31.89 \pm 3.11$ | $-133.39 \pm 14.77$ | $103.43 \pm 12.42$ |
| I222K | $-25.62 \pm 4.81$ | $-183.84 \pm 16.84$ | $-27.64 \pm 2.87$ | $211.48 \pm 15.95$ | $185.86 \pm 12.63$ |
| H274Y | $-24.98 \pm 4.37$ | $-139.67 \pm 12.33$ | $-24.91 \pm 2.60$ | $-164.59 \pm 10.99$ | $139.61 \pm 9.05$ |
| H274Y-I222K | $-19.70 \pm 3.30$ | $-82.20 \pm 8.85$ | $-28.72 \pm 2.59$ | $-110.92 \pm 8.28$ | $91.22 \pm 7.57$ |

$\Delta G_{\text{bind}}$ —binding free energy; $\Delta E_{\text{ele}}$ —electrostatic interaction; $\Delta E_{\text{vdw}}$ —van der Waals forces; $\Delta E_{\text{gas}}$ —gas-phase interaction; $\Delta G_{\text{sol}}$ —solvation energy

**Table 2.** Crystal structures and PDB codes, and abbreviation of simulated systems.

| Simulated System | PDB code | Abbreviation |
| --- | --- | --- |
| H5N1 wild type | 2HU4 | WT |
| H5N1 I222K | 2HU4 | I222K |
| H5N1 H274Y | 3CLO | H274Y |
| H274Y-I222K | 3CLO | H274Y-I222K |

These abbreviations are used throughout the text

In table 1, the wild type and I222K displayed a difference of −82.34 kcal/mol (ΔE_ele_) and −4.25 kcal/mol (ΔE_vdw_), respectively. Whilst H274Y and H274Y-I222K showed a difference of −57.47 kcal/mol (ΔE_ele_) and −3.81 kcal/mol (ΔE_vdw_), respectively. The difference in the intermolecular interacting components between the enzyme and ligand is demonstrative of a more stable system for the wild type and H274Y mutation in comparison to the I222K and H274Y-I222K mutation systems. The solvation energy ΔG_sol_, between the wild type and I222K, was calculated to have a difference of 82.43 kcal/mol, whilst that of H274Y and H274Y-I222K was found to be 48 kcal/mol. This difference was largely contributed to the change to a polar amino acid such as Ile to Lys and His to Tyr in the mutation systems. The estimated difference in binding free energy ΔG_bind_ for the wild type and I222k was calculated to be −4.34 kcal/mol. Whilst that of H274Y and H274Y-I222K was calculated to be −5.28 kcal/mol. The trend in binding free energy difference was observed to be as follows: WT_H5N1_<I222K_H5N1_<H274Y_H5N1_<H274Y-I222K_H5N1_, which indicates that the protein-ligand interaction became progressively thermodynamically unfavourable in the presence of different mutations. These calculations agree with experimental data that indicated that the H274Y-I222K mutation leads to a 10- to 90-fold increased relative resistance towards Peramivir [13]

### 2.5. Per-Residue Interaction Energy Decomposition Analysis

MM/GBSA was applied to gain a deeper understanding of the amino acid residues that are crucial for ligand-protein interactions. Figure 6 displays the energy decomposition of protein-ligand interactions per residue for wild type, I222K, H274Y, and H274Y-I222K mutant complexes.

**Figure 6.**
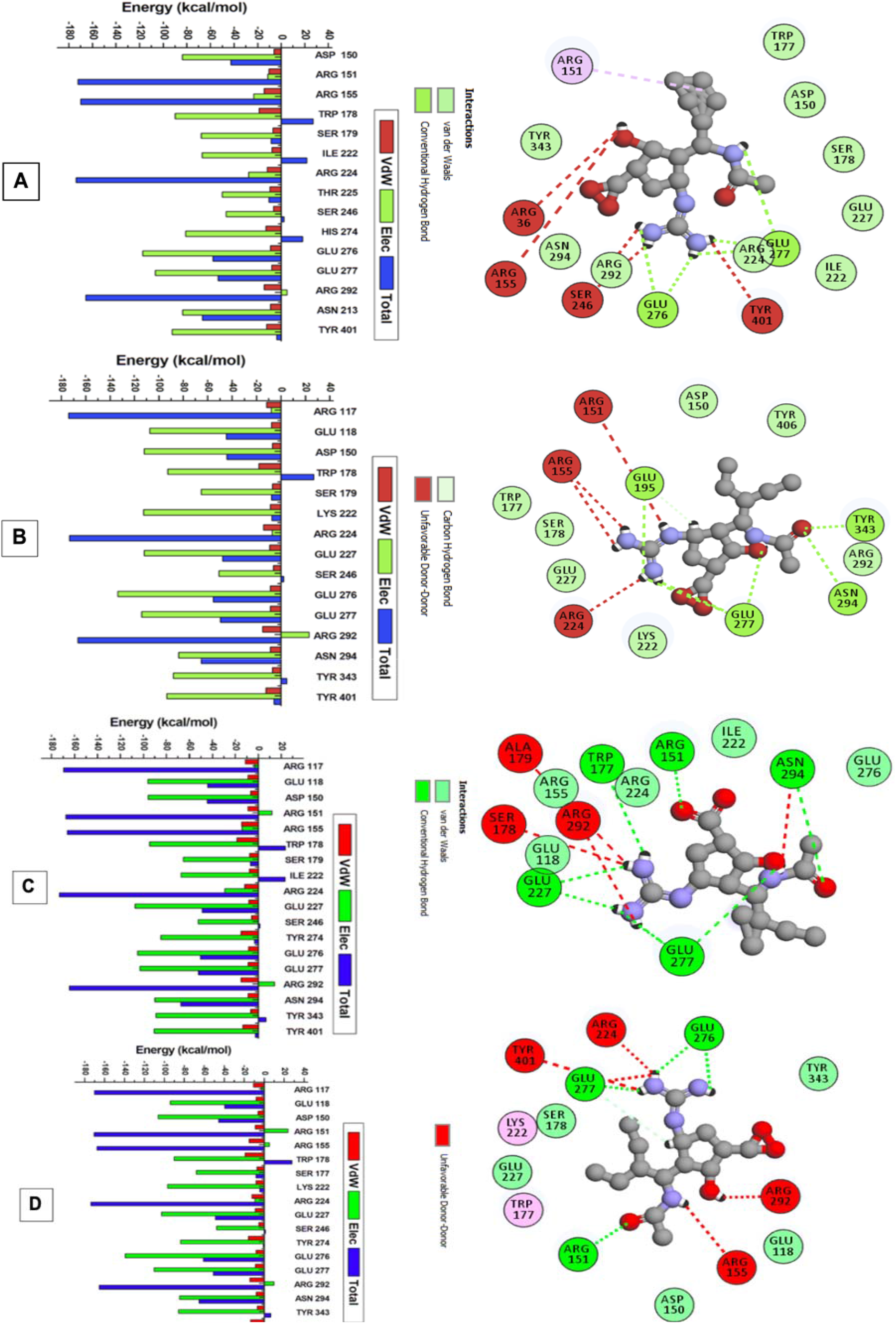
The per-residue energy decomposition analysis of Peramivir-wild type (A), Peramivir-I222K mutant (B), Peramivir-H274Y mutant (C), and Peramivir-H274Y-I222K mutant (D) complexes.

Figure 6(a, and b) showed that the wild type Ile222 complex had a larger residual energy contribution. As opposed to this, the mutant Lys222 complex contributed little energy to the Peramivir complex. As shown in Figure 6(a, and c), the wild type His247 complex exhibited a high residual energy contribution, whereas the mutant Tyr274 complex exhibited a low contribution to the residual energy of Peramivir. I222K and H274Y mutations may have affected the conformation of the protein-ligand complex at the binding site, resulting in this effect. Further destabilization of the protein-ligand complex was caused by the H274Y-I222K mutation, as shown in Figure 6d. The protein-ligand interaction at the active site (Figure 6d) also suggests that Lys222 has a weaker binding interaction compared to wild type and I222K. Based on these results, the protein-ligand interaction became progressively more unfavourable as different mutations were introduced.

The energy decomposition analysis revealed that Arg117, Arg224, and Arg292 contributed the largest residual energy to Peramivir binding to wild type, I222K, H274Y, and H274Y-I222K mutants, as shown in Figure 6. This is also in accordance with our previous study, which indicated that these residues contributed significantly to Peramivir binding to wild type and mutant H274K [18]. It is thought that these residues play a crucial role in forming the binding pocket between Peramivir and neuraminidase. The stronger interaction between protein and ligand, which occurs in the wild type complex, can be attributed to the overall higher level of total energy contribution and lower degree of flexibility of the backbone C-α atoms.

### 2.6. Principal Components Analysis (PCA)

PCA tool was applied to characterize the conformational variations between the wild type and I222K, H274Y and H274Y-I222K mutations. The two principal components (PC1 vs PC2) for all the systems were plotted for a comprehensive analysis of the conformational dynamics of neuraminidase structural enzymes [19]. Figure 7 demonstrates the variations of the amino acid residues of the wild type and I222K, H274Y and H274Y-I222K mutations.

**Figure 7.**
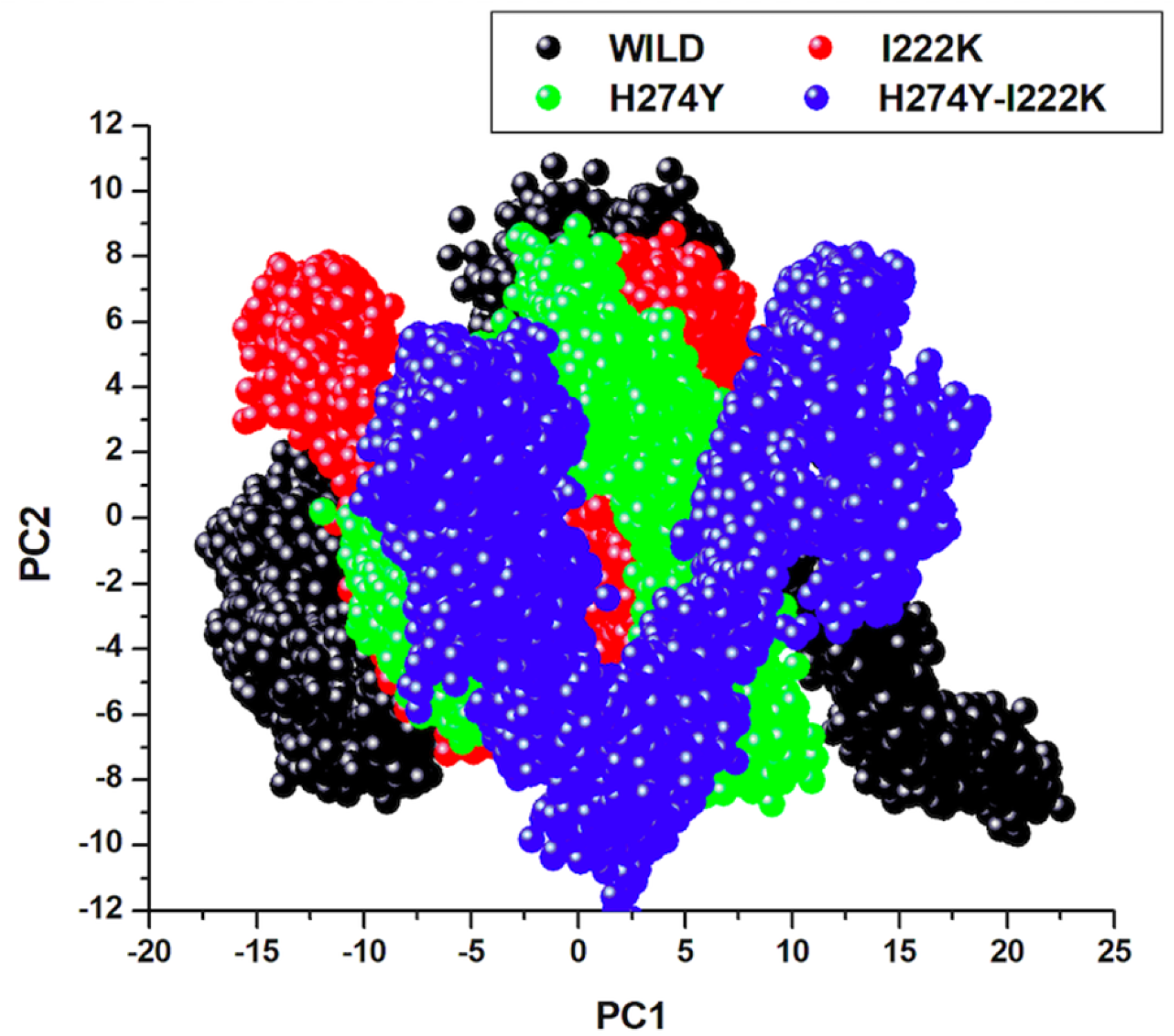
Projection of PC1 over PC2 for the wild type, I222K, H274Y and H274Y-I222K mutant systems measured over 200,000 ps MD simulation.

Figure 7 illustrates that the H274Y mutant occupies a larger space in comparison to the wild type. Therefore, this suggests that the wild type possesses a lower degree of flexibility, enabling the HRV enzyme to bind easily than the H274Y mutant. The H274Y mutant has an inconsistent residual motion with a correlation R^2^ value of 0.00055347, indicating a single cooperative direction, whilst the wild type has a correlation R^2^ value of 0.0017546, which shows a more proportionate motion. As evident in Figure 3 and Figure 4, the H274Y-mutant has more flexibility in comparison to wild type. From the evidence in Figure 7, a basic understanding of the dynamic behaviour of the biological systems is attained.

## 3. Materials and Methods

### 3.1. System Preparation

Wild type (H5N1) and mutant (H274Y) neuraminidase X-ray crystal structures were extracted from the RCSB Protein Data Bank (PDB: 2HU4), and (PDB: 3CL0), respectively [20]. Peramivir was docked into the active site using AutoDock Tools [21]. The proteins were prepared following the standard procedure using UCSF Chimera software package [22]. An introducing point mutation at positions 222 from Isoleucine (Ile) to Lysine (K) and 274 from Histidine (H) to Tyrosine (Y) to obtain the mutant structures using UCSF Chimera. Table 1 lists the four enzymes with the relevant mutations.

### 3.2. Molecular Docking

AutoDock Vina software was applied to calculate the docking scores with Geister partial charges were being allocated during the docking process. Lamarckian generic algorithm was applied to source docked conformations. The grid box size was x = 30 Å, y = 30 Å, z = 26 Å for the dimensions and x= −29.612 Å, y= −55.925 Å and z = 10.321 Å. The grid box houses the active site residues; ARG117, GLU 118, ASP 150, ARG 151, TRP 178, SER 179, GLU 200, ARG 224, GLU 146, ASP 151, ARG 152, ILE 222, SER 246, ALA 250, HIS 274, GLU 277, ASP 279, ARG 292, ASN 294 and TYR 401. All systems were docked with Peramivir.

### 3.3. Molecular Dynamic (MD) Simulations

MD simulations setup on the wild type and mutations, H274Y, I222K, H274Y-I222K, respectively using GPU AMBER14 software package [23–25]. In our previous work, system setup, minimizations, heating and equilibration steps were thoroughly explained [26–28].

### 3.4. Binding Energy Calculations

The binding free-energy profiles of Peramivir bound to H5N1 neuraminidase were computed using the Molecular Mechanics/Generalized Born Surface Area (MM/GBSA) approach [29–31]. From binding free energy, valuable insight into the association of the protein-ligand in a complex is obtained and is considered to be the end point energy calculation. The binding free energy was calculated taking into account 1000 snapshots from 200 000 ps trajectories.

The following set of equations describes the calculation of the binding free energy:

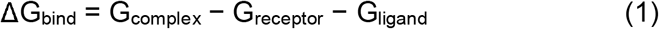

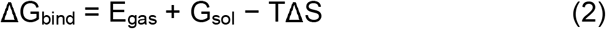

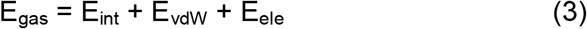

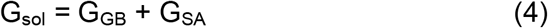

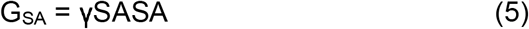

where E_gas_ denotes gas-phase energy; E_int_ signifies internal energy; and E_ele_ and E_vdw_ indicate the electrostatic and Van der Waals contributions, respectively. E_gas_ is the gas phase, elevated directly from the FF14SB force terms. G_sol_ denotes solvation free energy, can be decomposed into polar and nonpolar contribution states. The polar solvation contribution, G_GB_, is determined by solving the GB equation, whereas, G_SA_, the nonpolar solvation contribution is estimated from the solvent accessible surface area (SASA) determined using a water probe radius of 1.4 Å. T and S correspond to temperature and total solute entropy, respectively. To determine the contribution of each of the amino acids towards total binding free energy profile between H274Y, I222K and H274Y-I222K a per-residue decomposition analysis of the interaction energy for each residue was carried out using the MM/GBSA method.

### 3.5. Results Visualization and Analysis

UCSF chimera [32] was employed for visual analysis while statistical data and plots were analyzed using Microcal Origin analytical software (Origin Lab, Northampton, MA, http://www.originlab.com). A total continuous MD simulation time of 200 000 ps was performed, and the trajectories were then saved for every 1ps and analyzed. CPPTRAJ and PTRAJ modules [33] were used for analysis: Root Mean Square Deviation (RMSD), Root Mean Square Fluctuation (RMSF), Thermodynamic Calculation, and Principal Components Analysis (PCA).

### 3.6. Principal Components Analysis (PCA)

Principal components analysis (PCA) also referred to as essential dynamics (ED) and is one of the utmost unconventional methods for trajectory analysis. PCA has proved to be powerful and robust and opens new opportunities to visualize and explore the dynamics of protein cavities [34,35]. PCA describes the eigenvectors and eigenvalues, which represent the direction of motions and the amplitudes in those directions of the protein, respectively [36]. After stripping the ions and solvent of the 200 000 ps MD trajectories, PCA was performed on C-α atoms over 1000 snapshots at a time interval of 100 ps using the CPPTRAJ module in AMBER 14 in computing the first two principal components (PC1 and PC2), the corresponding PCA scatter plots were generated using Origin software.

## 4. Conclusions

To elucidate the versatile nature of the resistance of I222K, H274Y and H274Y-I222K mutation to Peramivir, we explored different computational strategies. Based on RMSD, RMSF, RoG, and PCA profiles, the Peramivir-wild type complex showed low residue flexibility and higher conformational stability. We observed the following trend in binding free energy difference: WT<I222K<H274Y<H274Y-I222K, indicating that the protein-ligand interaction became progressively thermodynamically unfavourable in the presence of different mutations. The introduction of H274Y and I222K mutations, therefore, affect Peramivir’s optimal orientation and conformation within the active site. A decrease in binding affinity results from the thermodynamic instability of mutant complexes, which results in weakened binding interactions, thus reducing Peramivir’s potency. In the wild type complex, the higher total energy contributions and low residue flexibility are thought to account for strong protein-ligand interactions. This study provides a premise for further investigation into the effects of other mutations on Peramivir’s efficacy against H5N1. In addition, we recommend combination therapy with approved influenza H5N1 virus drugs, as this can reduce the incidence of drug resistance. The results of this study promise to provide insights and knowledge that will enhance future drug designs and be helpful in the fight against the H5N1 virus.

## Institutional Review Board Statement

Not applicable.

## Informed Consent Statement

Not applicable.

## Data Availability Statement

The data presented in this study are available on request from the corresponding author.

## Acknowledgments

The authors acknowledge the UKZN School of Health Sciences and the Center for High-Performance Computing (CHPC, http://www.chpc.ac.za, accessed on 25 November 2021) for computational resources.

## References

1. Lai, S.; Qin, Y.; Cowling, B.J.; Ren, X.; Wardrop, N.A.; Gilbert, M.; Tsang, T.K.; Wu, P.; Feng, L.; Jiang, H.; et al. Global epidemiology of avian influenza A H5N1 virus infection in humans, 1997-2015: a systematic review of individual case data. Lancet. Infect. Dis. 2016, 16, e108–e118, doi:10.1016/S1473-3099(16)00153-5.

2. Farooqui, A.; Huang, L.; Wu, S.; Cai, Y.; Su, M.; Lin, P.; Chen, W.; Fang, X.; Zhang, L.; Liu, Y.; et al. Assessment of Antiviral Properties of Peramivir against H7N9 Avian Influenza Virus in an Experimental Mouse Model. Antimicrob. Agents Chemother. 2015, 59, 7255–7264, doi:10.1128/AAC.01885-15.

3. Lu, Y.; Li, Z.; Ma, C.; Wang, H.; Zheng, J.; Cui, L.; He, W. The interaction of influenza H5N1 viral hemagglutinin with sialic acid receptors leads to the activation of human γδ T cells. Cell. Mol. Immunol. 2013, 10, 463–470, doi:10.1038/cmi.2013.26.

4. Sauer, A.-K.; Liang, C.-H.; Stech, J.; Peeters, B.; Quéré, P.; Schwegmann-Wessels, C.; Wu, C.-Y.; Wong, C.-H.; Herrler, G. Characterization of the Sialic Acid Binding Activity of Influenza A Viruses Using Soluble Variants of the H7 and H9 Hemagglutinins. PLoS One 2014, 9, e89529.

5. Daudé, D.; Topham, C.M.; Remaud-Siméon, M.; André, I. Probing impact of active site residue mutations on stability and activity of Neisseria polysaccharea amylosucrase. Protein Sci. 2013, 22, 1754–1765, doi:10.1002/pro.2375.

6. Webster, R.G.; Peiris, M.; Chen, H.; Guan, Y. H5N1 outbreaks and enzootic influenza. Emerg. Infect. Dis. 2006, 12, 3–8, doi:10.3201/eid1201.051024.

7. Koh, G.; Wong, T.; Cheong, S.; Koh, D. Avian Influenza: a global threat needing a global solution. Asia Pac. Fam. Med. 2008, 7, 5, doi:10.1186/1447-056X-7-5.

8. Hsu, K.-C.; Hung, H.-C.; HuangFu, W.-C.; Sung, T.-Y.; Eight Lin, T.; Fang, M.-Y.; Chen, I.-J.; Pathak, N.; Hsu, J.T.-A.; Yang, J.-M. Identification of neuraminidase inhibitors against dual H274Y/I222R mutant strains. Sci. Rep. 2017, 7, 12336, doi:10.1038/s41598-017-12101-3.

9. Mtambo, S.E.; Amoako, D.G.; Somboro, A.M.; Agoni, C.; Lawal, M.M.; Gumede, N.S.; Khan, R.B.; Kumalo, H.M. Influenza Viruses: Harnessing the Crucial Role of the M2 Ion-Channel and Neuraminidase toward Inhibitor Design. Molecules 2021, 26, 880, doi:10.3390/molecules26040880.

10. Li, L.; Li, Y.; Zhang, L.; Hou, T. Theoretical studies on the susceptibility of oseltamivir against variants of 2009 A/H1N1 influenza neuraminidase. J. Chem. Inf. Model. 2012, 52, 2715–2729, doi:10.1021/CI300375K/SUPPL_FILE/CI300375K_SI_001.PDF.

11. Karthick, V.; Shanthi, V.; Rajasekaran, R.; Ramanathan, K. Exploring the Cause of Oseltamivir Resistance Against Mutant H274Y Neuraminidase by Molecular Simulation Approach. Appl. Biochem. Biotechnol. 2012, 167, 237–249, doi:10.1007/S12010-012-9687-7.

12. Wang, N.X.; Zheng, J.J. Computational studies of H5N1 influenza virus resistance to oseltamivir. Protein Sci. 2009, 18, 707, doi:10.1002/PRO.77.

13. Huang, L.; Cao, Y.; Zhou, J.; Qin, K.; Zhu, W.; Zhu, Y.; Yang, L.; Wang, D.; Wei, H.; Shu, Y. A Conformational Restriction in the Influenza A Virus Neuraminidase Binding Site by R152 Results in a Combinational Effect of I222T and H274Y on Oseltamivir Resistance. Antimicrob. Agents Chemother. 2014, 58, 1639, doi:10.1128/AAC.01848-13.

14. Hess, B. Convergence of sampling in protein simulations. Phys. Rev. E. Stat. Nonlin. Soft Matter Phys. 2002, 65, 31910, doi:10.1103/PhysRevE.65.031910.

15. Sneha, P.; George Priya Doss, C. Chapter Seven - Molecular Dynamics: New Frontier in Personalized Medicine. In Personalized Medicine; Donev, R.B.T.-A. in P.C. and S.B., Ed.; Academic Press, 2016; Vol. 102, pp. 181–224 ISBN 1876-1623.

16. Huang, Y.; Paul, D.R. Effect of molecular weight and temperature on physical aging of thin glassy poly(2,6-dimethyl-1,4-phenylene oxide) films. J. Polym. Sci. Part B Polym. Phys. 2007, 45, 1390–1398, doi:10.1002/polb.21173.

17. Pan, L.; Patterson, J.C. Molecular dynamics study of Zn(abeta) and Zn(abeta)2. PLoS One 2013, 8, e70681, doi:10.1371/journal.pone.0070681.

18. Buthelezi, N.M.; Mhlongo, N.N.; Amoako, D.G.; Somboro, A.M.; Sosibo, S.C.; Shunmugam, L.; Machaba, K.E.; Kumalo, H.M. Exploring the impact of H5N1 neuraminidase (H274Y) mutation on Peramivir: a bio-computational study from a molecular perspective. J. Biomol. Struct. Dyn. 2019, 38, 4344–4352, doi:10.1080/07391102.2019.1677501.

19. Principal Component Analysis and Factor Analysis BT - Principal Component Analysis. In; Jolliffe, I.T., Ed.; Springer New York: New York, NY, 2002; pp. 150–166 ISBN 978-0-387-22440-4.

20. Berman, H.M.; Westbrook, J.; Feng, Z.; Gilliland, G.; Bhat, T.N.; Weissig, H. The protein data bank. Nucleic Acids Res 2000, 28, doi:10.1093/nar/28.1.235.

21. Trott, O.; Olson, A.J. AutoDock Vina: improving the speed and accuracy of docking with a new scoring function, efficient optimization, and multithreading. J. Comput. Chem. 2010, 31, 455–461, doi:10.1002/jcc.21334.

22. Goddard, T.D.; Huang, C.C. FerrinTE. Software extensions to UCSF chimera for interactive visualization of large molecular assemblies. Structure 2005, 13, doi:10.1016/j.str.2005.01.006.

23. Go□ Tz, A.W.; Williamson, M.J.; Xu, D.; Poole, D.; Le Grand, S.; Walker, R.C. Routine Microsecond Molecular Dynamics Simulations with AMBER on GPUs. 1. Generalized Born. 2012, doi:10.1021/ct200909j.

24. Salomon-Ferrer, R.; Götz, A.W.; Poole, D.; Le Grand, S.; Walker, R.C. Routine Microsecond Molecular Dynamics Simulations with AMBER on GPUs. 2. Explicit Solvent Particle Mesh Ewald. J. Chem. Theory Comput. 2013, 9, 3878–3888, doi:10.1021/ct400314y.

25. Salomon-Ferrer, R.; Case, D.A.; Walker, R.C. An overview of the Amber biomolecular simulation package. Wiley Interdiscip. Rev. Comput. Mol. Sci. 2013, doi:10.1002/wcms.1121.

26. Bhakat, S.; Martin, A.J.M.; Soliman, M.E.S. An integrated molecular dynamics, principal component analysis and residue interaction network approach reveals the impact of M184V mutation on HIV reverse transcriptase resistance to lamivudine. Mol. Biosyst. 2014, 10, 2215–2228, doi:10.1039/c4mb00253a.

27. Chetty, S.; Soliman, M.E.S. Possible allosteric binding site on Gyrase B, a key target for novel anti-TB drugs: homology modelling and binding site identification using molecular dynamics simulation and binding free energy calculations. Med. Chem. Res. 2015, 24, 2055–2074, doi:10.1007/s00044-014-1279-3.

28. Kumalo, H.M.; Soliman, M.E. A comparative molecular dynamics study on BACE1 and BACE2 flap flexibility. J. Recept. Signal Transduct. 2016, 36, 505–514, doi:10.3109/10799893.2015.1130058.

29. Kollman, P.A.; Massova, I.; Reyes, C.; Kuhn, B.; Huo, S.; Chong, L.; Lee, M.; Lee, T.; Duan, Y.; Wang, W.; et al. Calculating structures and free energies of complex molecules: combining molecular mechanics and continuum models. Acc. Chem. Res. 2000, 33, 889–897.

30. Massova, I.; Kollman, P.A. Combined molecular mechanical and continuum solvent approach (MM-PBSA/GBSA) to predict ligand binding. Perspect. Drug Discov. Des. 2000, 18, 113–135, doi:10.1023/A:1008763014207.

31. Tsui, V.; Case, D.A. Theory and applications of the generalized Born solvation model in macromolecular simulations. Biopolymers 2000, 56, 275–291, doi:10.1002/1097-0282(2000)56:4<275::AID-BIP10024>3.0.CO;2-E.

32. Pettersen, E.F.; Goddard, T.D.; Huang, C.C.; Couch, G.S.; Greenblatt, D.M.; Meng, E.C.; Ferrin, T.E. UCSF Chimera--a visualization system for exploratory research and analysis. J. Comput. Chem. 2004, 25, 1605–12, doi:10.1002/jcc.20084.

33. Roe, D.R.; Cheatham, T.E. PTRAJ and CPPTRAJ: Software for Processing and Analysis of Molecular Dynamics Trajectory Data. J. Chem. Theory Comput. 2013, 9, 3084–3095, doi:10.1021/ct400341p.

34. Desdouits, N.; Nilges, M.; Blondel, A. Principal Component Analysis reveals correlation of cavities evolution and functional motions in proteins. J. Mol. Graph. Model. 2015, 55, 13–24, doi:10.1016/j.jmgm.2014.10.011.

35. Martínez, L. Automatic Identification of Mobile and Rigid Substructures in Molecular Dynamics Simulations and Fractional Structural Fluctuation Analysis. PLoS One 2015, 10, e0119264, doi:10.1371/journal.pone.0119264.

36. Cocco, S.; Monasson, R.; Weigt, M. From Principal Component to Direct Coupling Analysis of Coevolution in Proteins: Low-Eigenvalue Modes are Needed for Structure Prediction. PLoS Comput Biol 2013, 9, e1003176.

